# α-Synuclein facilitates endocytosis by elevating the steady-state levels of phosphatidylinositol 4,5-bisphosphate

**DOI:** 10.1101/2020.06.18.158709

**Authors:** Schechter Meir, Atias Merav, Abd Elhadi Suaad, Davidi Dana, Gitler Daniel, Sharon Ronit

**Author notes:** **Corresponding author:** Ronit Sharon, PhD.

## Abstract

α-Synuclein (α-Syn) is a protein implicated in the pathogenesis of Parkinson’s disease (PD). It is an intrinsically disordered protein that binds acidic phospholipids. Growing evidence supports a role for α-Syn in membrane trafficking, including, mechanisms of endocytosis and exocytosis, although the exact role of α-Syn in these mechanisms is currently unclear. Here we have investigated the role of α-Syn in membrane trafficking through its association with acidic phosphoinositides (PIPs), such as phosphatidylinositol 4,5-bisphosphate (PI4,5P_2_) and phosphatidylinositol 3,4-bisphosphate (PI3,4P_2_). Our results show that α-Syn colocalizes with PIP_2_ and the phosphorylated active form of the clathrin adaptor AP2 at clathrin-coated pits. Using endocytosis of transferrin, an indicator of clathrin mediated endocytosis (CME), we find that α-Syn involvement in endocytosis is specifically mediated through PI4,5P_2_ levels. We further show that the rate of synaptic vesicle (SV) endocytosis is differentially affected by α-Syn mutations. In accord with their effects on PI4,5P_2_ levels at the plasma membrane, the PD associated E46K and A53T mutations further enhance SV endocytosis. However, neither A30P mutation, nor Lysine to Glutamic acid substitutions at the KTKEGV repeat domain of α-Syn, that interfere with phospholipid binding, affect SV endocytosis. This study provides evidence for a critical involvement of PIPs in α-Syn-mediated membrane trafficking.

**Significance Statement:** α-Synuclein (α-Syn) protein is known for its causative role in Parkinson’s disease. α-Syn is normally involved in mechanisms of membrane trafficking, including endocytosis, exocytosis and synaptic vesicles cycling. However, a certain degree of controversy regarding the exact role of α-Syn in these mechanisms persists. Here we show that α-Syn acts to increase plasma membrane levels PI4,5P_2_ and PI3,4P_2_ to facilitate clathrin mediated and synaptic vesicles endocytosis. Based on the results, we suggest that α-Syn interactions with the acidic phosphoinositides facilitate a shift in their homeostasis to support endocytosis.

## Introduction

α-Synuclein (α-Syn) protein is critically implicated in the pathogenesis of Parkinson’s disease (PD). α-Syn reversibly interacts with membrane lipids. It preferentially binds acidic phospholipids (Davidson et al., 1998; Stockl et al., 2008; Bodner et al., 2009) and fatty acids (Sharon et al., 2001, 2003a, 2003b). Upon lipid binding, the intrinsically disordered α-Syn protein acquires a α-helical structure (Davidson et al., 1998).

α-Syn plays a role in membrane trafficking and synaptic vesicle cycling (Lautenschläger et al., 2017), however, its exact role in these mechanisms is far from being clear. In a previous study, we reported the first indication for a role of α-Syn in clathrin-mediated endocytosis (CME) and synaptic vesicle (SV) cycling (Ben Gedalya et al., 2009). We suggested that α-Syn acts to increase membrane curvature through enrichment of membrane phospholipids with poly-unsaturated fatty acids (PUFAs) and increased membrane fluidity (Sharon et al., 2003b; Ben Gedalya et al., 2009). It was further suggested based on α, β, γ-Syn knock-out in mice that all three synucleins are involved in clathrin-mediated synaptic vesicle (SV) recycling at presynaptic nerve terminals (Vargas et al., 2014).

Side by side with studies reporting an activating role, other studies reported an inhibitory role for α-Syn in endocytosis. Experiments performed in mice expressing α-Syn at the calyx of Held indicated an inhibitory role in presynaptic endocytosis (Xu et al., 2016). α-Syn was shown to inhibit SV endocytosis during intense electrical stimulation of lamprey neurons (Busch et al., 2014). Furthermore, neurons overexpressing α-Syn internalized lower amounts of styryl dyes, which serve as indicators for SV recycling, suggesting a reduction in endocytosis (Scott et al., 2010; Scott and Roy, 2012).

In addition to its reported role(s) in endocytosis, a large body of evidence indicates a role for α-Syn in mechanisms of exocytosis. These include the soluble NSF attachment proteins (SNAP) receptor (SNARE) complex assembly (Burre et al., 2010; Darios et al., 2010; Thayanidhi et al., 2010) or SNARE protein binding (Sun et al., 2019) and vesicle fusion (Choi et al., 2013; DeWitt and Rhoades, 2013; Lai et al., 2014); transmitter release (Larsen et al., 2006; Lundblad et al., 2012); and the regulation of fusion pore dilation (Logan et al., 2017). However, a certain degree of controversy regarding the exact role for α-Syn in exocytosis persists.

Phosphoinositides (PIP)s are acidic membrane phospholipids that control many aspects of cell biology. The most common form in the plasma membrane is PI4,5P_2_, which among other cellular functions, regulates mechanisms of endocytosis and exocytosis. In endocytosis, PI4,5P_2_ is involved in the early steps of recruiting endocytic clathrin adaptors and their accessory factors to the plasma membrane (recently reviewed in (Kaksonen and Roux, 2018)). Its enrichment with PUFAs (Balla, 2013) may contribute to the formation or maintenance of membrane curvature, which is critical for the initiation of a clathrin coated pit in endocytosis (Avinoam et al., 2015; Haucke and Kozlov, 2018). With maturation of the endocytic vesicle, PI4,5P_2_ is converted to PI3,4P_2_ to allow completion of the process (Posor et al., 2013; He et al., 2017). In exocytosis, PI4,5P_2_ is involved in priming and fusion steps of Ca^2+^-triggered vesicle release (Martin, 2015). PI4,5P_2_ recruits and activates specific proteins that regulate SNARE complex assembly and function (Martin, 2012, 2015). Additional roles for PI4,5P_2_ in dilation of the fusion pore have been described (Martin, 2012, 2015). These may involve PI4,5P_2_-binding proteins such as CAPS and synaptotagmin (Martens et al., 2007; Lynch et al., 2008; Hui et al., 2009) or its effects on F-actin polymerization (Berberian et al., 2009).

In this study, we report that α-Syn’s involvement in mechanisms of transferrin and SV endocytosis is specifically mediated through its activity to enrich the plasma membrane with PI4,5P_2_ and PI3,4P_2_ (PIP_2_). Our results point at PIP_2_ as key components in α-Syn-mediated mechanisms of membrane trafficking in neuronal and non-neuronal cells.

## Methods

### Plasmids

WT α-Syn or the specified α-Syn mutations were cloned into a pcDNA vector (Zarbiv et al., 2014). Costume-ready Mission shNIR2 (TRCN0000029763); shSNCA (TRCN0000272292) and shCntrl were from Sigma-Aldrich (Rehovot, Israel) (Schechter et al., 2020). The following plasmids were used: pGFP-C1-PLCδ1-PH (Addgene # 21179 from Tobias Meyer (Stauffer et al., 1998)); CFP-FKBP-Ins4,5p and Lyn-FRB (Suh et al., 2006); BFP_2_-INPP4B-CAAX (Goulden et al., 2019); and INPP5E-mCherry (Posor et al., 2013). In addition, an adeno-virus expressing Synaptophysin-2XpHluorin under the human synapsin (hSyn) promotor (AAV1/2 hSyn:Synaptophysin-2XpHluorin) and AAV1/2 hSyn:WT α-Syn (Atias et al., 2019) were used. AAV1/2 hSyn:mutant α-Syn were constructed by PCR amplification followed by complete sequencing of the inserts.

### Cells

HEK 293T, HeLa, SH-SY5Y and an inducible α-Syn-expressing SH-SY5Y cell line (Davidi et al., 2020) were maintained in Dulbecco's modified eagle's medium (DMEM) supplemented with 10% FBS; 2% L-glutamine; 1% penicillin/streptomycin, sodium-pyruvate and non-essential amino acids (Biological Industries, Beit-Haemek, Israel). α-Syn expression was induced in the inducible SH-SY5Y cell line with 1 μM/ml doxycycline (Sigma-Aldrich, Rehovot, Israel). SK-Mel2 cells express detectable levels of endogenous α-Syn, however, these are lowered with passages. Thus, a large number of aliquots at passage twelve were kept frozen and experiments were performed between weeks 2-6 from thawing a frozen aliquot. SK-Mel2 cells were maintained in minimum essential medium (MEM; Sigma-Aldrich, Rehovot, Israel) supplemented with 10% FBS; 1% L-glutamine, penicillin/streptomycin and sodium-pyruvate. Cultures were maintained at 37°C in a 95% air/ 5% CO_2_ humidified incubator.

### Primary cultures

Primary hippocampal cultures were prepared by dissecting both hippocampi of P0-P1 C57BL/6JOlaHsd (α-Syn^−/−^) mouse brains, as described previously (Gitler, 2004; Orenbuch et al., 2012). After trituration in 20 units/ml papain solution (Worthington Lakewood, NJ, USA), 1×10^5^ cells were plated on coverslips that were pre-coated with 5 μg/ml poly-D-lysine (Sigma-Aldrich, Rehovot, Israel) in a 24-well plate containing 1.5 ml Neurobasal-A medium (Gibco, Thermo Fisher Scientific, Petah Tikva, Israel) supplemented with 5% fetal bovine serum, 2% B-27 (Gibco, Thermo Fisher Scientific), 1% Glutamax I, and 1μg/ml gentamicin. After 1 day, the solution was replaced to 1 ml Neurobasal-A supplemented with 2% B-27 and 1% Glutamax I. To slow the proliferation of glial cells, 1 μM cytosine β-D-arabinofuranoside (Ara-C; Sigma-Aldrich) was added to the culture at 2 days in vitro (DIV). Cultures were maintained at 37°C in a 5% CO_2_ humidified incubator until used at 13 DIV.

### Synaptophysin-pHluorin (sypHy) imaging

Neurons were infected at 5 DIV with AAV1/2 hSyn:sypHy and were imaged at 13 DIV. Coverslips were placed in a field stimulation chamber (Warner Scientific, Hamden, CT, USA) in an extracellular solution composed of the following (in mm): 150 NaCl, 3 KCl, 20 glucose, 10 HEPES, 2 CaCl_2_, 2 MgCl_2_, pH adjusted to 7.35 with NaOH at 310 mOsm. The solution also contained glutamate receptor antagonists APV (50μM) and DNQX (10μM) to avoid recurrent network activity. Neurons were imaged at room temperature every 6 seconds on a Nikon TiE inverted microscope equipped with a Neo5.5 Andor sCMOS camera, using an EGFP filter set (Chroma, Bellows Falls, VT, USA). After acquiring 6 baseline images (*F*_0_), neurons were stimulated by applying 300 bipolar pulses at 20 Hz, each of 1 ms duration and 10V/cm amplitude, through parallel platinum wires. At the completion of the experiment, the culture was exposed to saline in which 50 mM of the NaCl_2_ was replaced with NH4Cl, to expose the total pool of vesicles (Fmax; (Atias et al., 2019)). The background-corrected fluorescence values recorded for each synapse were normalized either by the peak response during the stimulation train, or by the size of the total pool of vesicles, as indicated. The rate of endocytosis was assessed by exponential fitting of the time course of the decay in fluorescence from its peak upon the completion of stimulation back to baseline values. To exclusively image exocytosis, we added 1 μM bafilomycin-A (BafA) to the extracellular solution. BafA blocks the vesicle proton pump, thus masking the endocytotic segment of SV cycle (Burrone et al., 2006) without affecting the kinetics of endocytosis (Qiu et al., 2015). Quantification was performed with the NIS-Elements software (Nikon), by placing equal circular regions of interests (ROI)s on 30-50 synapses in each field and extracting the background-subtracted average fluorescence value of each ROI (Atias et al., 2019). A local background was obtained adjacently to each ROI.

### Viral production and transduction

AAV1/2 particles were produced as previously described (Orenbuch et al., 2012). Briefly, HEK293T cells were co-transfected with the pD1 and pD2 helper plasmids and a plasmid containing the cDNA of interest located between AAV2 ITRs, preceded by the hSyn promotor. After 3 days of incubation at 37 °C in a humidified 5% CO_2_ incubator, cells were lysed in lysis solution (150 mM NaCl, 50 mM Tris-HCl pH 8.5) using 3 rapid freeze-thaw cycles (in an ethanol bath chilled to −80 °C and a heated 37°C water bath). The supernatant was treated with 10 units/ml benzonaze (Sigma-Aldrich, Rehovot, Israel), cleared by centrifugation and filtrated through a 0.45 μm membrane. The viral particles were maintained at 4°C until use. Viral titer was determined by infecting neuronal cultures, aiming for 80-90% infection efficiency, verified by immunofluorescence or direct fluorescence imaging, as applicable. Viral titer was determined by adding 0.2-2 μl of the viral prep directly to the growth medium at 5 DIV.

Lentiviral particles were produced as described (Schechter et al., 2020) by co-transfecting HEK 293T cells with a set of three plasmids: pCMVΔR8.91; pMD2.G; and a transfer plasmid pLKO-1-puro. Virus titer was determined for each preparation following transduction of cells, by quantitative PCR using specific primers for Puromycin resistance gene: forward, 5’-TCACCGAGCTGCAAGAACTCT-3’ and reverse primer, 5’-CCCACACCTTGCCGATGT-3’. Primer sequence for human SNCA: forward: 5’-GCAGGGAGCATTGCAGCAGC-3 and reverse 5’-GGCTTCAGGTTCGTAGTCTTG-3’; Nir2: forward: 5’-GCTTTGATGCACTCTGCCAC-3’ and reverse: 5’-AGCTCATTGTTCATGCTCCC-3’; G6PD: forward: 5’-CACCATCTGGTGGCTGTTC-3 and reverse 5’-TCACTCTGTTTGCGGATGTC-3; SK-Mel2 and SH-SY5Y cells were infected by incubating the cells (1× 10^6^) in FBS free-DMEM, containing viral particles and polybreane (4μg/ml) for 6 hours. The conditioning medium was then replaced with 10% FBS-supplemented DMEM.

### FACS

Analysis was performed as previously described (Schechter et al., 2020). Briefly, cells were fixed in 2% paraformaldehyde at 4 °C and permeabilized in 0.2% saponin in 1% BSA (w/v) for 15 minutes at 4°C. Cells were then incubated with anti α-Syn antibody (MJFR1, 1:2,000 Abcam, Zotal, Tel Aviv, Israel) and anti PIP abs (Echelon Biosciences, Salt Lake City, UT), including: PI3P (Z-P003; 1:300), PI4P (Z-P004 1:300), PI3,4P_2_ (Z-P034b;1:300), PI4,5P_2_ (Z-A045; 1:200) and PI3,4,5P_3_ (Z-P345b; 1:400), for 90 minutes with gentle rolling; washed and probed with the respective secondary antibody for 30 minutes at room temperature. Analyses were performed using BD LSRFortessa Cell Analyzer, equipped with 5 lasers (355, 405, 488, 561 and 640 nm) and the FLOWJO, LLC software. Each experiment also included relevant compensation controls. A control consisting of cells grown and processed in parallel, treated with 10 μM ionomycin (Balla, 2013) for 5-10 minutes at room temperature, was also included. Gating was based on FSC, SSC and positive immunoreactivity for α-Syn. A total of 2000-4000 gated cells were counted in each experiment unless indicated differently.

### Transferrin endocytosis

Measurement of transferrin endocytosis were performed as previously described (Ben Gedalya et al., 2009; Posor et al., 2013) with some modifications. Cells were grown in 12-wells plates, on cover slides that were pre-treated with poly-D-lysine (100 μg/ml) for 1 hour. On the day of the experiment, cells were serum-starved for 1.5 hours. Cells were then conditioned in 25 μg/ml of 568-Transferrin (568-Tf; Molecular Probes, Invitrogen, Rhenium, Israel) in clear DMEM for 7 min at 37 °C. When specified, induction of FRB-FKBP dimerization and recruitment of Inp45p to the PM was achieved with the addition of rapamycin (500 nM) in DMSO (0.5% v/v). After two washes with ice-cold PBS, cells were acid washed at pH 5.3 (0.2 M sodium acetate, 0.2 M sodium chloride) on ice for 1.5 min, to remove surface-bound transferrin. Cells were then washed 2 additional times with ice-cold PBS, fixed in 2% paraformaldehyde (PFA) for 20 min on ice and processed for ICC.

### PI4,5P_2_ detection by the PH-PLCδ1-GFP biosensor

Cells were grown in 12-well plates, on cover slides that were pre-coated with poly-D-Lysine (100 μg/ml, for 1 hour). Cells were co- transfected to express PH-PLCδ1-GFP together with WT α-Syn or one of the specified α-Syn mutations (A30P, E46K, A53T, K10,12E or K21,23E), or a mock plasmid. In some experiments, cells were conditioned in the presence of 50 μg/ml 647 Concanavalin A (ConA, molecular probes, Invitrogen, Rehovot, Israel) in DMEM, at 37 °C for 10 minutes to label the plasma membrane. Membranes were defined by the ring-shaped ConA signal around the cell and were differentiated from the cytoplasm. Membrane to cytosolic PH-PLCδ1-GFP signal ratio was calculated using the NIS-Element AR Analysis 4.20.02 64-bit software (Nikon, Agentek, Tel Aviv, Israel)(Schechter et al., 2020).

### Immunocytochemistry (ICC)

Cell lines or primary neuronal cultures were fixed in cold 2% PFA for 20 minutes, washed in PBS and permeabilized with 0.5% saponin in blocking solution (1% BSA in PBS w/v) for 30 minutes at room temperature. Cells were incubated with the following anti α-Syn antibodies, C20 (Santa Cruz, Dallas TX, US) at 1:500 dilution, overnight at 4°C; MJFR-1 (Abcam, Zotal, Tel Aviv, Israel; 1:2000), anti-α-Syn (ab21976 1:330, Abcam); P-AP2M1-T156 (D4F3; Cell Signaling Technology; 1:300). The following antibodies were from Echelon Biosciences (Salt Lake City, UT, USA): anti PI3,4P_2_ (Z-P034b;1:300) or anti PI4,5P_2_ (Z-A045; 1:800) for 2 hours at room temperature. Cells were then washed (PBS; 10 min x 3) and then incubated with a host-suitable secondary ab, washed again, and mounted in vectashield mounting medium (Vector-labs, Burlingame, CA USA).

### Fluorescence microscopy and image analysis

Images were acquired using a Zeiss LSM 710 Axio Observer confocal Z1 laser scanning microscope, equipped with an argon laser 488, Diode 405-430 laser and HeNe 633 laser.

For colocalization analyses, images were captured using Nikon’s A1R+ confocal microscope, equipped with an ultrahigh-speed resonant scanner and high-resolution digital galvano scanner, with four laser unit LU-N4S. Per each experiment, the exciting laser, intensity, background levels, photo multiplier tube (PMT) gain, contrast and electronic zoom were maintained constant. The antibody-specific background was subtracted. The focus of each picture was obtained by choosing the plane with greatest fluorescent signal. Quantifications were performed with NIS-element software. A constant threshold for the signal of α-Syn and each PIP was maintained to all images. The nucleus was excluded from quantification. The program automatically defined the positive spots for either α-Syn or each PIP. Then, it automatically calculated the number of positive pixels in colocalization for each channel. Results were normalized to total α-Syn positive pixels.

### Western Blotting

Samples of lysed cells (70 μg protein) were loaded on a 10% SDS-PAGE and following electrophoresis were transferred to a PVDF membrane. Membrane was blocked with milk (10% in TBST) and reacted with anti Nir2 ab (ab22823, Abcam, 1:500). The immuno-blots were reacted with a secondary, HRP-conjugated, anti-goat antibody (1:100,000, Jackson ImmunoResearch), visualized with Clarity Western ECL Substrate (BioRad, Rishon Le Zion, Israel) and scanned with ChemiDoc XRS+ imaging system (Bio-Rad). Signal density was quantified using UN-SCAN-IT GEL 3.1 software (Silk Scientific, Orem, UT, USA).

### Detection of P-Ser129 α-Syn

The detection of PSer129 α-Syn levels was performed as described previously (Davidi et al., 2020). Briefly, Cells were lysed in homogenization buffer [20 mM Hepes, pH 7.4; 1 mM MgCl2; 0.32 M sucrose; 43 mM 2-mercaptoethanol; 1 x protease inhibitors] by eight passages through a 27-gauge needle. The homogenate was then centrifuged at 10 000 x g for 10 min, at 4°C. The resultant supernatant was analyzed by Lipid-ELISA (Abd-Elhadi et al., 2016). Samples (0.5 μg protein) were loaded onto a 96-well white PolySorp ELISA plate, that was precoated with phospholipids and GM-1 ganglioside. Plates were incubated for 3 hours at 37°C to allow capture of proteins by the immobilized lipids. Lipid-bound PSer129 α-Syn was detected with anti PSer129 α-Syn ab (pSyn#64, WAKO, Osaka, Japan) diluted 1:1000 in 1% BSA in PBS followed by incubation with an HRP-conjugated secondary ab. Detection reaction with Super Signal ELISA Femto (Pierce, Ornat, Israel), luminescence was determined by Luminometer (Infinite M200 Pro. NEOTEL Scientific Instrumentation Ltd.) immediately after adding the substrate. A standard curve consisting of recombinant protein, phosphorylated at Ser129 (Proteos, Kalamazoo, US), was applied in parallel to the test samples and used as a reference.

PSer129 α-Syn levels were normalized to total α-Syn levels, detected in the same sample with anti α Syn ab MJFR 1 (Abcam) HRP-conjugated. Detection reaction with TMB one component microwell substrate (Southern Biotech, Birmingham, Alabama, USA) – 100 μl well. The reaction was terminated with 100 μl/well of 1M H_2_SO_4_. Absorbance at 450 nm was determined using a plate-reader (EL808 Ultra Microplate Reader, Bio-Tek Instruments, VT, USA). Standard curve of purified human α-Syn protein was used for reference.

## Experimental Design and Statistical Analysis

All experiments were performed in parallel with their designed controls and in random order, and they were replicated at least three times. Data are shown as mean ± SE. Statistical comparisons were performed with the two-tailed Student's t-test. When multiple comparisons were performed, we applied the Bonferroni correction. The distribution of the variables in each experimental group was normal. All tests were conducted using Statistical Package for the Social Sciences (SPSS) version 20.0. Significant differences were accepted at P < 0.05.

## Results

### α-Syn colocalizes with phosphorylated AP2 (pAP2) and PIP_2_ on clathrin coated pits (CCP)

Phosphorylation at Thr156 of μ2 subunit of the clathrin adaptor AP2 starts following its binding to PI4,5P_2_ at the initiation of a CCP and throughout vesicle lifetime (Ricotta et al., 2002; Kadlecova et al., 2017; Wrobel et al., 2019). We analyzed SK-Mel2 cells, which express detectable levels of endogenous α-Syn protein. The immunoreactive signal for pAP2, observed by ICC, appeared on the plasma membrane and colocalized with α-Syn (Fig. 1A). To assess the specificity of pAP2 signal we utilized a specific inhibitor (LP-935509) for Numb-associated kinases (NAKs), that phosphorylate the μ2 subunit of AP2 (Ricotta et al., 2002; Kostich et al., 2016; Wrobel et al., 2019). pAP2 signal was dramatically reduced in cells treated with the LP-935509 inhibitor (10 μM, for 3 hours) and no obvious colocalization of α-Syn and pAP2 could be detected (Fig. 1B). The specificity of α-Syn signal was confirmed in cells that their α-Syn expression was silenced with shSNCA and treated with the LP-935509 inhibitor. The results show a substantial loss of both signals, α-Syn and pAP2 (Fig. 1C). To confirm that α-Syn colocalizes with pAP2 on CCP, the slides were immunoreacted also with antibodies against PI4,5P_2_ (Fig. 1) or PI3,4P_2_, which regulates the initiation steps and maturation of the clathrin coated vesicle (Posor et al., 2013; He et al., 2017) (Supplementary Fig. 1). The results show a substantial degree of colocalization for the immunoreactive signals obtained for α-Syn, pAP2 and either PI4,5P_2_ or PI3,4P_2_.

**Fig. 1.**
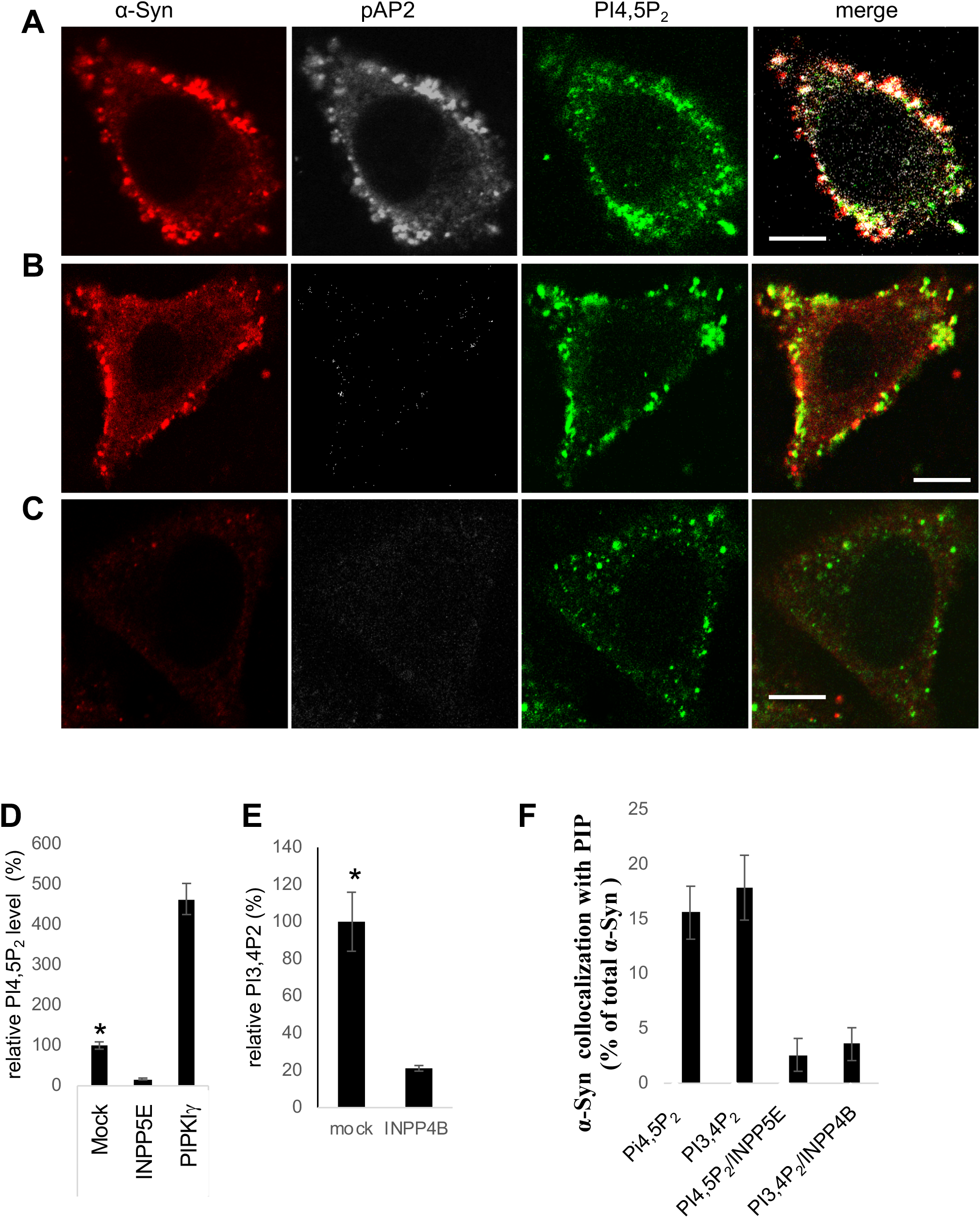
α-Syn colocalizes with phosphorylated AP2 and PIP_2_ on clathrin-coated pits. **A.** SK-Mel2 cells were processed for the detection of the immunoreactive signals of endogenous α-Syn (MJFR1 ab, red), phosphorylated Thr153 μ2 subunit of AP2 (pAP2; gray) and PI4,5P_2_ (green) by immunocytochemistry (ICC). Bar= 10 μM. **B**. SK-Mel2 cells treated with LP-935509 (10 μM, for 3 hours), to inhibit the phosphorylation of the μ2 subunit of AP2 and processed for ICC as in (A). Bar= 10 μM. **C**. SK-Mel2 cells infected with lentivirus encoding shSNCA to silence α-Syn expression. Cells were treated with LP-935509 inhibitor as in (B) and processed for ICC. Bar= 10 μM. **D**. HEK 293T cells, transfected to express the specified PIP-metabolizing proteins. Cells analyzed by FACS to immunodetect PI4,5P_2_ levels. N>2000 cells in each group; mean ± SE * P<0.01, t-test. **E**. Cells analyzed by FACS as in (D) to detect PI3,4P_2_ signal. N>2000 cells; mean ± SE * P<0.01, t-test **F.** SK-Mel2 cells processed for ICC as in (A) and co-immunoreacted with anti α-Syn and anti PIP Abs (Echelon). Colocalization of the signal obtained for α-Syn with the specified PIP was quantified and normalized to total α-Syn positive spots. Colocalization is reduced in controls cells that express the specific PIP-phosphatase. N>25 cells; mean ± SE.

The specificity of PI4,5P_2_ and PI3,4P_2_ signals were assessed in control SK-Mel2 cells, transfected to express either the inositol polyphosphate-5-phosphatase E (INPP5E) or the inositol polyphosphate-4-phosphatase B (INPP4B), that dephosphorylate the phosphate at the 5- or −4 position of the inositol ring, respectively; or type Iγ PI4P-5-kinase (PIPKIγ), that produces PI4,5P_2_. Importantly, PI4,5P_2_ signal was dramatically lower in cells expressing the INPP5E phosphatase, and ~4.5 fold higher with the expression of PIPKIγ (Fig. 1D). In accord, PI3,4P_2_ signal was dramatically lower with the expression of INPP4B phosphatase (Fig. 1E; FACS; n> 2,000 cells in each group; P<0.01; t-test).

Using a program-based method, we scanned the ICC images to identify positive pixels in each channel and the colocalizing pixels within the channels. A portion of α-Syn signal specifically colocalized with PI4,5P_2_ (~16%) and PI3,4P_2_ (~18%). Colocalization of α-Syn with PI4,5P_2_ or PI3,4P_2_ was diminished following the expression of INPP5E or INPP4B, respectively (Fig. 1F; mean ± SE, n>25 cells). These results suggest that endogenous α-Syn localizes, at least in part, to PI4,5P_2_ / PI3,4P_2_-positive endocytic clathrin-coated pits, consistent with a possible function in CME.

### α-Syn involvement in endocytosis of transferrin associates with alterations in PIP_2_ levels

Endocytosis of fluorescently labelled transferrin was utilized as a functional readout for CME. Endogenous α-Syn expression was downregulated in the SK-Mel2 cells using shSNCA and were 37% of the levels detected in control cells, infected with shCntrl (Figs. 1C and 2A-B). α-Syn levels were kept down regulated for at least 14 days and experiments were performed during this time window. Endocytosis of 568-Tf was significantly (58%) lower in shSNCA than in shCntrl cells (set at 100%, Fig. 2A, B). In agreement with our recent report (Schechter et al., 2020), silencing α-Syn expression resulted in significantly lower levels of PI4,5P_2_ (73%) compared with control cells (set at 100%). Similarly, PI3,4P_2_ levels were also lower (75%) with silencing α-Syn expression (ICC, n=17-25 cells in each group; t-test; P<0.05).

**Fig. 2:**
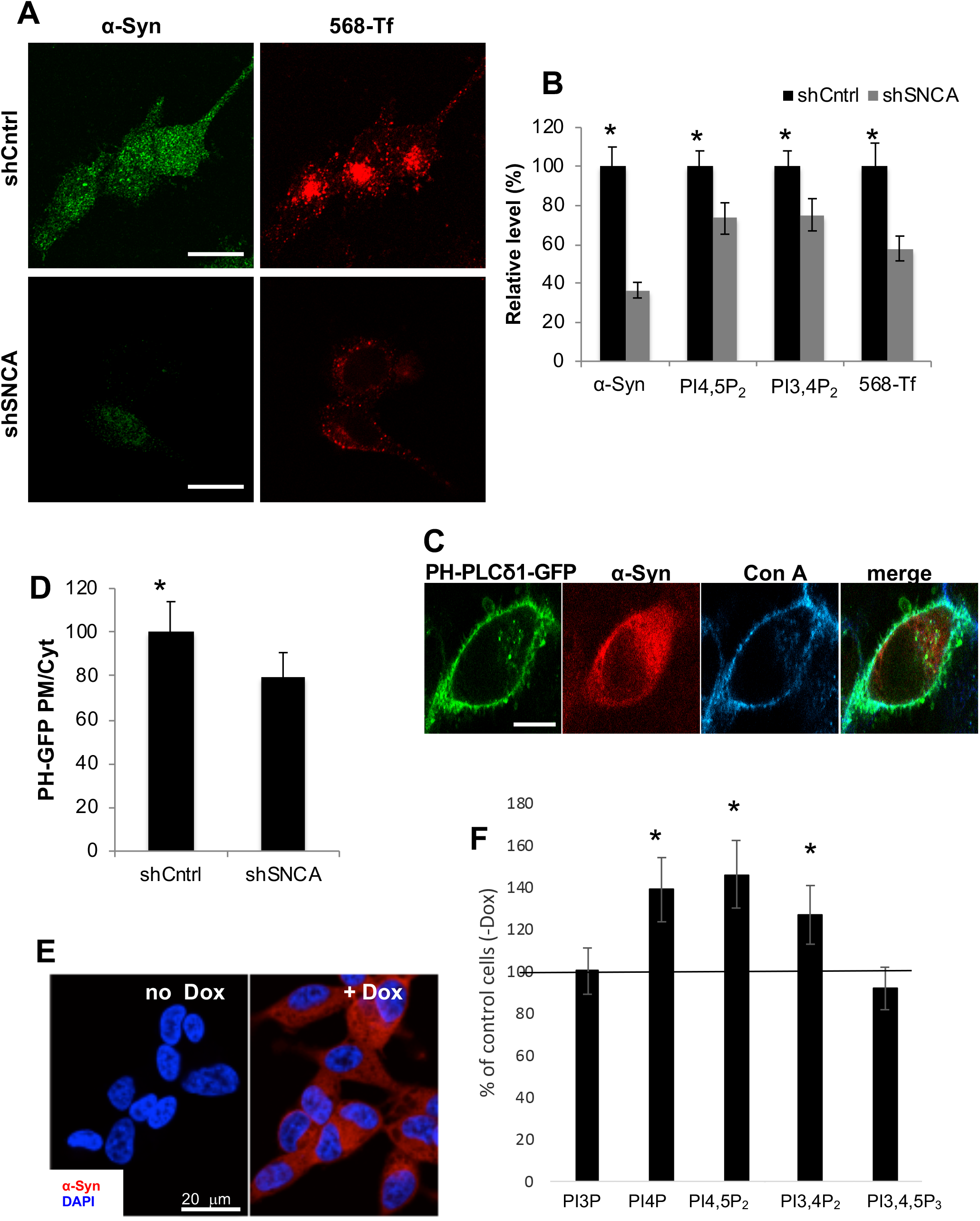
α-Syn increases PIP_2_ levels and endocytosis of 568-Tf. **A**. SK-Mel2 cells infected with lentivirus encoding shSNCA or shCntrl were processed for endocytosis of 568-Tf (red) and detection of α-Syn with an anti-α-Syn ab (MJFR1; green). Bar=20μm. **B**. SK-Mel2 cells expressing shSNCA or shCntrl (as in (A)). The immunoreactive signals for α-Syn, PI4,5P_2_, PI3,4P_2_ and 568-Tf detected by ICC (mean ± SE of n= 15-22 cells per treatment;*, P<0.05 t-test) **C**. Cells expressing PH-PLCδ1-GFP were incubated with 647-ConA to mark the plasma membrane, fixed and immunoreacted with anti α-Syn ab (C20; red). The direct fluorescence of the GFP (Green) and 647-Con A (Blue) is demonstrated. Bars =20μm. **D.** The PH-PLCδ1-GFP signal ratio (plasma membrane to cytosol) determined in SK-Mel2 cells expressing shCntrl or shSNCA (n>15 cells per treatment, mean±SE; P<0.05 t-test). **E**. Inducible, Tet-on SH-SY5Y α-Syn-expressing cells incubated with doxycycline (1μg/ml) for 72h or with the DMSO solvent. Cells processed for ICC and immunoreacted with anti α-Syn ab (MJFR1; Red). DAPI staining depicts nuclei (blue). Bars= 20μm. **F.** α-Syn expression was induced with doxycycline (1μg/ml) for 72h in the inducible SH-SY5Y cells. Control cells were treated in parallel with the solvent. Cells immunoreacted with anti-PIP ab (Echelon) as specified, and analyzed by FACS. Results presented as percent of control cells, without induced α-Syn expression, set at 100%. N>2000 cells per treatment; mean ± SE; * P<0.05; t-test with Bonferroni correction for multiple comparisons.

To verify that the observed loss of PI4,5P_2_ was specific to the plasma membrane, we utilized the PH(PLCδ)-GFP biosensor for PI4,5P_2_ detection (Stauffer et al., 1998). SK-Mel2 cells expressing either shSNCA or shCntrl were transfected to express PH(PLCδ)-GFP. The signal ratio of GFP fluorescence in the plasma membrane, assessed with ConA (see methods) to cytosolic GFP was calculated and used to indicate plasma membrane PI4,5P_2_. Importantly, the results obtained with the PH(PLCδ)-GFP biosensor were highly similar to the results with anti-PI4,5P_2_ antibody (Fig. 2C,D) and confirmed the significant reduction in plasma membrane PI4,5P_2_ in α-Syn-depleted cells (i.e. 79% of PH(PLCδ)-GFP signal compared to control cells, set at 100%).

To assess the general effects of α-Syn on PIPs, we utilized an inducible SH-SY5Y cell line, expressing α-Syn under the control of doxycycline (Dox; Fig. 2E,F) (Davidi et al., 2020). α-Syn expression was induced for 72 hours and cells were processed for the detection of PIPs by FACS. Control cells that express a mock plasmid were treated in parallel. Significantly higher levels of PI4P, PI3,4P_2_ and PI4,5P_2_ were detected with inducing the expression of α-Syn compared with the control cells (set at 100%). In contrast, PI3P and PI3,4,5P3 levels were not altered upon α-Syn overexpression (Fig. 2F, mean ±SE, n>2000 cells, P<0.05. Data analyzed by t-test with Bonferroni correction for multiple comparisons).

These data suggest that α-Syn regulates the levels of PI3,4P_2_ and PI4,5P_2_ phosphoinositides that control CME of transferrin. We therefore decided to test the hypothesis that α-Syn increases PIP_2_ levels to enhance CME.

### α-Syn-mediated endocytosis of transferrin is PI4,5P_2_-dependent

To experimentally regulate the levels of PI4,5P_2_, we utilized an inducible enzymatic system to acutely deplete PI4,5P_2_ from the plasma membrane (Suh et al., 2006). This system enables rapamycin-induced targeting of Inp45p, a PI4P-5-phosphatase, to the plasma membrane. HEK293T cells were transfected to co-express the inducible phosphatase together either with WT α-Syn or pcDNA mock plasmid. 48 hours post DNA-transfection, cells were processed simultaneously for 568-Tf endocytosis together with activation of the phosphatase with rapamycin (see methods).

The Inp45p phosphatase is recruited to the plasma membrane in cells treated with rapamycin but remains in the cytoplasm in DMSO-treated cells (Fig. 3A). In accord, PI4,5P_2_ levels were lower in rapamycin (43%) compared with the DMSO treated cells (set at 100%), demonstrating phosphatase activity (ICC; Fig. 3B; P<0.01; t-test). To find out whether plasma membrane PI4,5P_2_ levels play a role in α-Syn’ effect to enhance endocytosis, we quantified 568-Tf internalization in cells that co-express Inp45p together with WT α-Syn and treated with rapamycin or DMSO (Fig. 3A, C). The results show that rapamycin-induced depletion of PI4,5P_2_ completely abolished the ability of overexpressed α-Syn to stimulate CME (Fig. 3C), whereas CME was stimulated by α-Syn expression in the DMSO-treated cells, measuring a higher degree of 568-Tf internalization (163%) compared with the control cells that express a mock pcDNA (set at 100%). In control cultures, in which cells were transfected and treated in parallel but without Inp45p expression, we found no effect for rapamycin on α-Syn-induced CME (Fig. 3C). Mean ± SE of n=4 experiments. *, P< 0.05. Data analyzed by t-test with Bonferroni correction for multiple comparisons.

**Figure 3:**
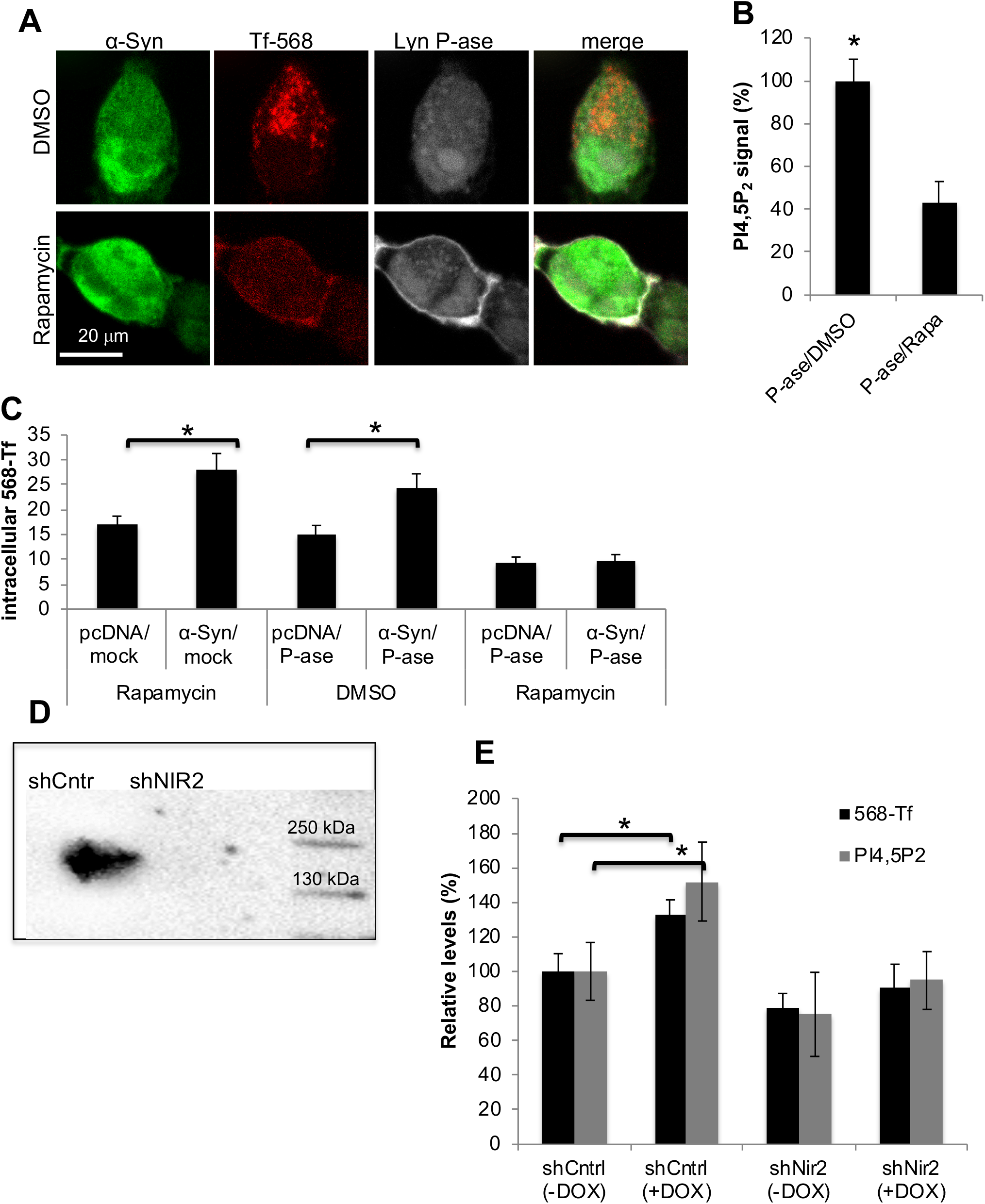
The enhancing effect of α-Syn on CME is PI4,5P_2_ dependent. **A.** HEK 293T cells, transfected to co-express the three plasmids: Lyn-FRB, FKBP-CFP-Inp54p and WT α-Syn. On day of experiment, cells were serum starved for 90 minutes and then incubated for 7 minutes with 568-Tf and rapamycin (500 nM) to induce the recruitment of the Inp45p phosphatase to the plasma membrane along with internalization of 568-Tf. Control cells were treated in parallel with DMSO. Cells were then acid washed, fixed and immunoreacted with anti α-Syn ab (MJFR1; Green). Direct fluorescence of CFP (Grey) and AF568-Tf (Red) was captured. **B.** HEK 293T cells expressing Lyn-FRB and FKBP-CFP-Inp54p were treated either with rapamycin or DMSO as described in (A), followed by immunoreaction with anti PI4,5P_2_ ab (Echelon). A quantification of PI4,5P_2_ signal is shown (n>15 cells per treatment; mean ± SE; * P <0.01 t-test. **C.** Graph showing quantification of internalized 568-Tf signal in cells transfected and treated as in (A). Mean ±SE of n>15 cells per treatment; * P <0.05 t-test; with Bonferroni correction for multiple comparisons. **D**. The inducible Tet-on SH-SY5Y were infected with shNir2 or a control shRNA (shCntrl). Protein samples were analyzed by a Western blot, immunoreacted with anti-Nir2 ab (Abcam). **E.** Cells as in (D) were induced to express α-Syn with doxycycline (1μg/ml, for 72h) or treated with an equivalent amount of DMSO solvent. Cells were then processed for ICC to measure 568-Tf endocytosis, as described in (C). Mean ± SE, n=20-25 cells per treatment; * P <0.05, t-test. Sister cultures were immunoreacted with anti PI4,5P_2_ ab and analyzed by FACS (n>2000 cells per treatment; mean ± SE; * P <0.05 t-test; with Bonferroni correction for multiple comparisons.).

To confirm a role of PI4,5P_2_ in α-Syn-mediated endocytosis of Tf-568, we tested the significance of silencing Nir2 expression. An important function of Nir2 protein is the exchange of endoplasmic reticulum phosphatidylinositol (PI) with PM phosphatidic acid (PA), which is required for maintaining PM levels of PI4,5P_2_ (Kim et al., 2013). Nir2 expression was silenced with shNir2 in the inducible α-Syn expressing SH-SY5Y cell line, resulting in lower Nir2 mRNA levels (~25%) and lower protein levels (~55%) of the respective levels detected in control cells, infected to express shCntrl (set at 100%, Fig 3D; P<0.01; t-test). α-Syn expression was then induced with doxycycline for 72 hours and cells were analyzed by FACS (n>2000 cells) to detect PI4,5P_2_ and α-Syn levels. Sister cultures were analyzed by ICC (n=20-25 cells) to detect internalized 568-Tf (Fig. 3E). Inducing α-Syn expression with doxycycline resulted in significantly higher PI4,5P_2_ (152%) levels and in accord, higher internalization of 568-Tf (133%), compared with cells that were not treated to induce the expression of α-Syn (100%). However, in cells that their Nir2 expression was silenced, induction of α-Syn expression had no effect on PI4,5P_2_ levels, nor endocytosis of 568-Tf. Setting the levels of PI4,5P_2_ and 568-Tf at 100% in cells induced to express α-Syn and infected to express the control shRNA, we detected 32% lower PI4,5P_2_ and 38% lower 568-Tf signals in cells treated with shNir. The results therefore show that silencing Nir2 significantly interfered with the effects of α-Syn to enhance the levels of PI4,5P_2_ and endocytosis of transferrin. Mean ±SE of 2-3 experiments, n>2000 cells in each treatment. *, P< 0.05, Data analyzed by t-test with Bonferroni correction for multiple comparisons.

### α-Syn mutations correlate endocytosis of transferrin with changes in plasma membrane levels of PI4,5P_2_

We tested three PD-associated mutations in α-Syn, A30P, E46K and A53T, and two synthetic K10,12E and K21,23E α-Syn mutation. The synthetic mutations were generated by replacing two positively charged Lysine residues within the KTKEGV repeat domain, with negatively charged Glutamic acid residues. In a previous study, these K to E mutations were shown to interfere with α-Syn binding to membrane phospholipids (Zarbiv et al., 2014).

Endocytosis of 568-Tf and PIP_2_ levels were determined in HEK 293T cells, expressing WT α-Syn or the specified α-Syn mutations. PI3,4P_2_ and PI4,5P_2_ levels were determined by FACS, using specific abs (n<2000 cells; Fig. 4) and plasma membrane levels of PI4,5P_2_ were determined by the PH(PLCδ)-GFP signal ratio (as above; n= 20-25 cells; Fig. 4). The results show significantly higher levels of 568-Tf endocytosis, PI3,4P_2_ and PI4,5P_2_ in WT α-Syn than in the mock-plasmid expressing cells. Further increases over WT α-Syn were generally detected for these measurables with the PD-associated mutations, with the exception of the A53T effect on PI3,4P_2_ and the E46K effect on plasma membrane PI4,5P_2_ levels (marked in red). The K-to-E mutations in α-Syn were not different from the mock-expressing cells (Fig. 4; mean ± SE Data analyzed by t-test with Bonferroni correction for multiple comparisons; P <0.05.

**Figure 4:**
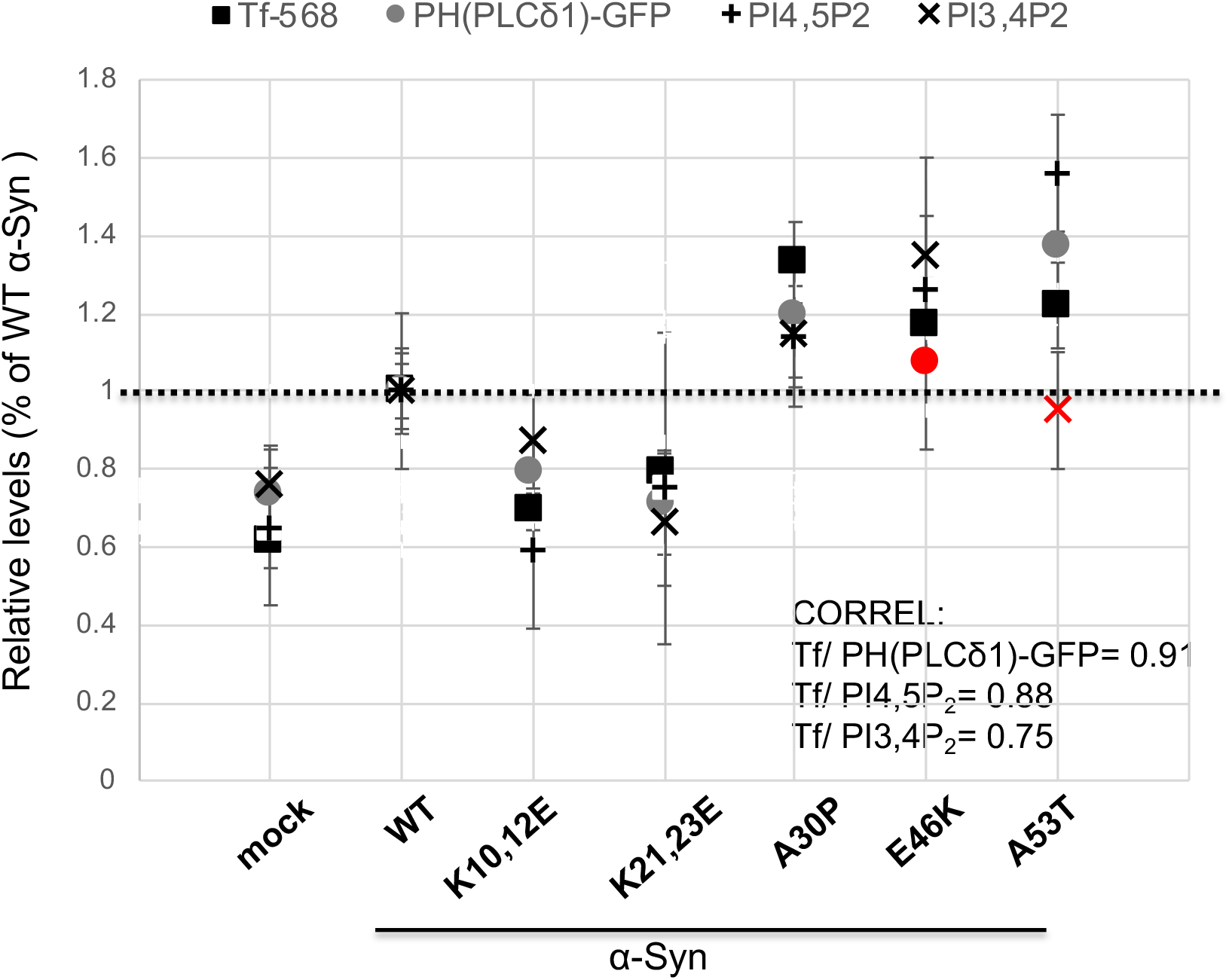
α-Syn mutations correlate endocytosis of transferrin with changes in plasma membrane levels of PI4,5P_2_. HEK 293T cells were transfected to express the indicated α-Syn forms. Cells were processed for 568-Tf endocytosis by ICC (n>15 cells per treatment). Sister cultures were immunoreacted with anti-PI4,5P_2_ ab or anti-PI3,4P_2_ ab and analyzed by FACS (n>2000 cells per treatment). Plasma membrane PI4,5P_2_ determined by calculating the membrane to cytosolic signal ratio of PH-PLCδ1-GFP in cells expressing WT α-Syn or the specified α-Syn mutations. 647-ConA was used to mark the plasma membrane. Vertical line represents the mock expressing cells. Mean ± SE of n=20-25 cells. Black marks indicate significance, P<0.05, t-test with Bonferroni correction. Red marks indicate statistical insignificance. CORREL, correlation coefficient.

A strong correlation between α-Syn effects on 568-Tf endocytosis and plasma membrane levels of PI4,5P_2_ was noted (correlation coefficient [r] = 0.91). Similarly, 568-Tf endocytosis correlated with total signal of PI4,5P_2_ (r = 0.88). 568-Tf endocytosis correlated also with PI3,4P_2_ levels (r = 0.75). We thus concluded that α-Syn increases PIP_2_ levels to facilitate CME and decided to test the hypothesis that it similarly acts to enhance SV endocytosis.

### α-Syn accelerates the rate of SV endocytosis alongside with reducing the fraction of released SVs

The involvement of α-Syn in SVs cycling was tested using Synaptophysin-pHluorin (sypHy) (Sankaranarayanan et al., 2000; Burrone et al., 2006). The pH-dependent fluorescence of sypHy, which is quenched in intact acidified SVs, increases upon exocytosis. After endocytosis and re-acidification of the SV lumen, fluorescence is re-quenched and returns to baseline. Primary hippocampal neurons prepared from α-Syn^−/−^ (C57BL/6JOlaHsd) mouse brains were infected to express sypHy together with one of the following α-Syn forms, WT α-Syn, the A30P, E46K, A53T mutants, or the two K to E mutations. mCherry served as a control for infection efficacy. SV cycling was measured at 13 DIV by imaging sypHy before, during and after the delivery of 300 stimuli at 20 Hz (Vargas et al., 2014; Atias et al., 2019). NH_4_Cl saline was applied following the return of fluorescence to baseline, to alkalinize all intracellular compartments, thus exposing the total size of the SV pool (F_max_).

The results show that WT α-Syn expression over the α-Syn^−/−^ background inhibits the extent of SV cycling, represented by a lower peak fluorescence (F_peak_/F_max_; Fig. 5A; n=50 synapses per image, 3 experiments. The lower peak level of the sypHy signal is in agreement with previous reports on an inhibitory role for α-Syn in exocytosis (Lautenschläger et al., 2017). A lower sypHy signal may arise either from a reduction in the number of SVs available for release, an acceleration of endocytosis or both. To assess specifically SV exocytosis, we added BafA to the bath. BafA inhibits re-acidification of the SVs after endocytosis and thus, sypHy measurements performed in its presence report exclusively exocytosis (Burrone et al., 2006). Indeed, in the presence of BafA, the cumulative sypHy signal in WT α-Syn expressing neurons was lower than in control neurons (Fig. 5B, C), indicating a reduction in the total secretory capacity of the presynaptic terminals, as has been previously reported (Nemani et al., 2010; Atias et al., 2019). Importantly, normalizing the traces by the peak fluorescence obtained at the completion of stimulation (DF/Fpeak) revealed that the kinetics of the decay of sypHy was accelerated by the expression of WT α-Syn (Fig. 5D, F). Thus, in addition to its inhibitory effect on the exocytotic segment of the SV cycle, α-Syn also accelerates the rate of endocytosis.

**Figure 5:**
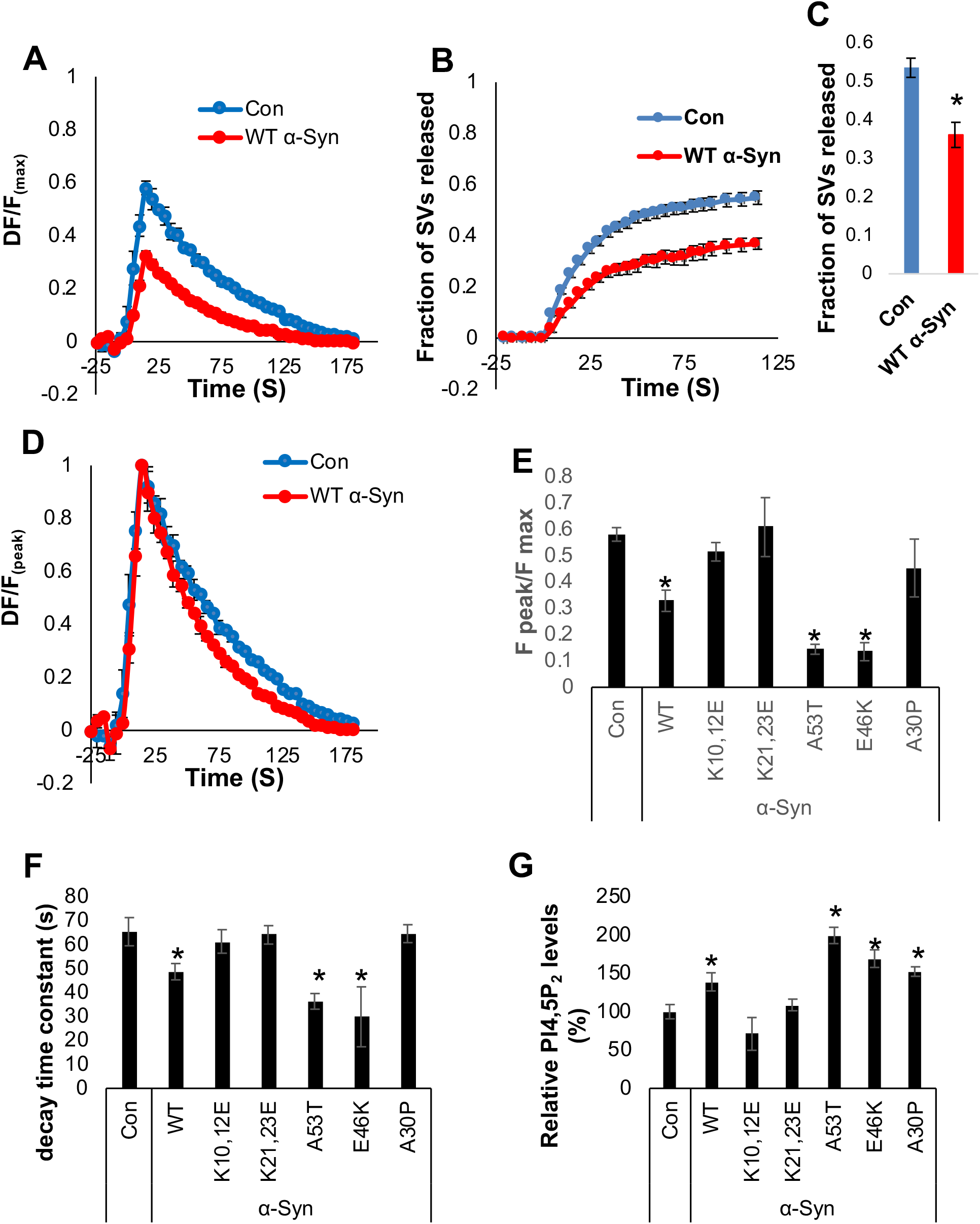
α-Syn mutations differentially affects SV endocytosis and PI4,5P_2_ levels. **A**. Hippocampal neurons at 13 DIV, prepared from α-Syn^−/−^ mouse brains were infected to express sypHy and mCherry, and either WT α-Syn or a mock plasmid. Neurons were stimulated for 15 seconds at 20Hz (300 stimuli) at room temperature and the change in sypHy fluorescence in the synaptic puncta was recorded. The baseline fluorescence prior to stimulation (F0) was subtracted. Fluorescence was normalized to the total pool of vesicles (F_max_) measured at the end of the measurements by exposure to NH_4_Cl-saline. Mean ± SE of n=6 slides per treatment (50 synapses per image, in 3 independent experiments). **B.** As in (A), but the bath included 10 μM BafA, and 2400 stimuli were applied at 20 Hz. **C**. Shown is the fluorescence measured 120 seconds after starting stimulation, normalized by the total pool of vesicles (F_max_), mean ± SE; n=12-20 slides, 30-50 synapses per image). P<0.001 t-test. **D**. Cells as in (A). Shown is ΔF scaled to the peak fluorescence (F_peak_). Mean ± SE; n=6 slides per treatment (50 synapses per image, in 3 independent experiments). **E.** Fractional peak release values (F_peak_/F_max_) for each of the specified α-Syn form. Shown are mean ± SE values; n=3-6 slides per treatment (30-50 synapses per image, 3 experiments). * P <0.05 t-test; with Bonferroni correction for multiple comparisons. **F.** Graph showing the calculated decay constant resembling the rate of endocytosis with each of the specified α-Syn form or the control plasmid. N=3-6 slides per treatment (30-50 synapses per image, 3 experiments).* P <0.05 t-test; with Bonferroni correction for multiple comparisons. **G.** Hippocampal neurons expressing the indicated α-Syn forms or a mock plasmid were processed for ICC at 13 DIV and immunoreacted with anti α-Syn ab (MJFR1) and anti PI4,5P_2_ ab (Echelon). Graph showing the quantification of PI4,5P_2_ in α-Syn positive neurites. Mean ± SE; N=8-13 fields per treatment * P <0.05 t-test; with Bonferroni correction for multiple comparisons.

The PD-associated mutations in α-Syn, E46K and A53T, further inhibited SV recycling (Fig. 5E) and in accord, further accelerated the rate of endocytosis (Fig. 5F). However, the A30P mutation and both K to E mutations were not different from control cells in their effects on SV recycling (Fig. 5 E,F). Together, measurements of α-Syn effects of SVs cycling, as determined by sypHy, reveal its complex effects on SV pools and architecture. However, considering the actual segment of SVs trafficking, α-Syn appears to accelerate the rate of endocytosis.

We next assessed PI4,5P_2_ levels in primary neurons, infected to express WT α-Syn or the specified mutations as above. At 13 DIV, neurons were fixed and processed for ICC with anti α-Syn and anti PI4,5P_2_ antibodies. Similar to the results in cell lines (Fig. 4) we found that expression of α-Syn mutations in hippocampal neurons differentially affected PI4,5P_2_ levels (Fig. 5G). That is, WT α-Syn increased PI4,5P_2_ levels (139%) over the levels detected in control cells (set at 100%); the PD-associated A30P, E46K and A53T mutations further increased PI4,5P_2_ levels (152-200%), however, PI4,5P_2_ levels in primary neurons expressing the K to E mutations did not differ from control cells. Mean±SE of n=8-13 fields. Data analyzed by t-test with Bonferroni correction for multiple comparisons; significance, P <0.05.

The results demonstrate a correlation between α-Syn-dependent increases in PI4,5P_2_ levels and its capacity to enhance the rate of endocytosis. That is, an inverse correlation of r= −0.75 was calculated between the decay-constant of sypHy signal and PI4,5P_2_ levels with the different α-Syn mutations. Excluding A30P mutation, which appears ineffective in endocytosis, yet increases PI4,5P_2_ levels, results in a stronger correlation (r= −0.87).

### Increases in cellular PI4,5P_2_ levels are associated with α-Syn toxicity

We next determined PSer129 α-Syn levels as an indication of α-Syn toxicity in HEK293T cells transfected to co-express α-Syn together either with INPP5E phosphatase or PIPKIγ kinase. PI4,5P_2_ levels were considerably lower with INPP5E and higher with PIPKIγ expression (Fig. 1D). Total and PSer129 α-Syn levels were determined in the soluble fraction of sister cultures by Lipid ELISA (see methods). PSer129 to total α-Syn ratio was calculated and found to be significantly higher in cells harboring high PI4,5P_2_ levels upon the expression of PIPKIγ (Fig. 6).

**Figure 6:**
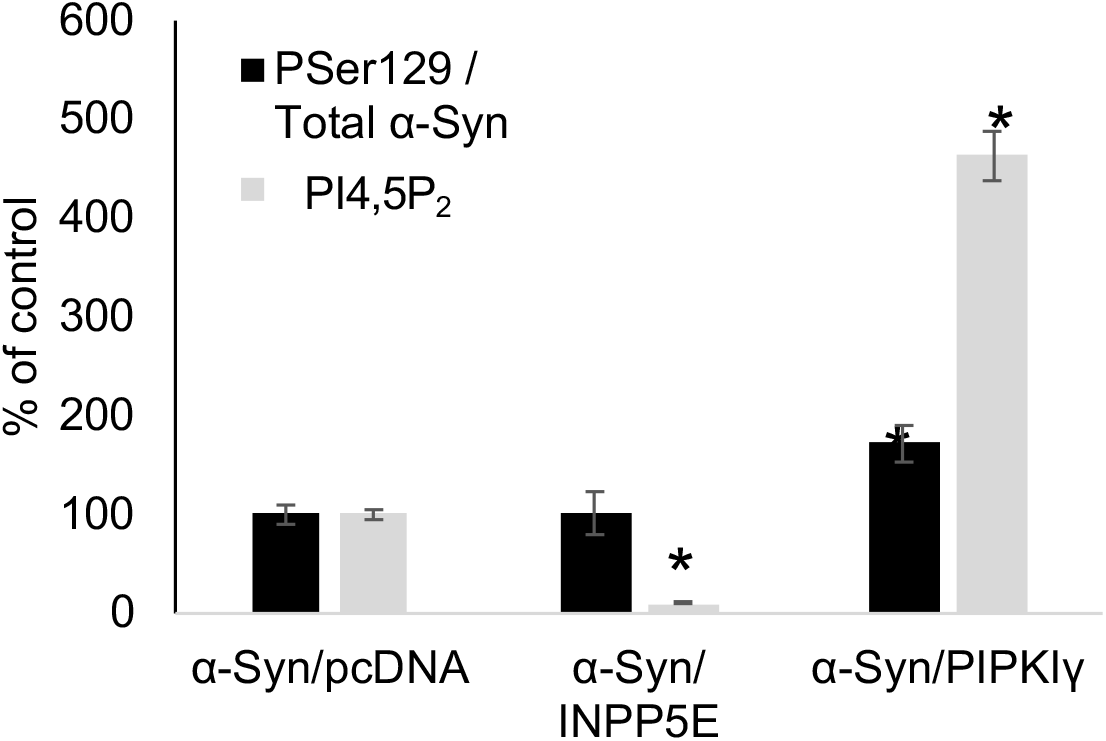
Increases in cellular PI4,5P_2_ levels are associated with α-Syn toxicity. HEK293T cells were transfected to express WT α-Syn together either with PIPKIγ or INPP5E. Control cells expressed WT α-Syn together with an empty pcDNA vector. PSer129 α-Syn levels were determined by phospholipids ELISA using anti PSer129 α-Syn ab and were normalized to total α-Syn levels detected in the same sample. Mean ±SE of n=2 independent experiments, each consisting of a triplicate. *, P<0.05

## Discussion

We address the controversy over α-Syn’s role in membrane trafficking and SV cycling by investigating α-Syn associations with PIP_2_ and specifically with PI4,5P_2_. We show that α-Syn colocalizes with components of clathrin coated pits/vesicles, e.g., pAP2, PI4,5P_2_ and PI3,4P_2_. We further show that α-Syn-mediated CME requires PI4,5P_2_ at the plasma membrane. Utilizing a recruitable 5-phosphatase, that hydrolyses PI4,5P_2_ at the plasma membrane, we demonstrate that α-Syn mediated CME is PI4,5P_2_-dependent. In addition, lowering PI4,5P_2_ levels by means of silencing the PI-transfer protein, Nir2, abolished the enhancing effect of α-Syn on 568-Tf endocytosis. Finally, utilizing specific α-Syn mutations, with differential effects on PI4,5P_2_ levels, we demonstrate a correlation between neuronal PI4,5P_2_ levels and the rate of SVs endocytosis, assessed by acidification of sypHy. Based on these results and the established role of PI4,5P_2_ in CME (Cremona et al., 1999; Haucke, 2005; Di Paolo and De Camilli, 2006; Zoncu et al., 2007; Posor et al., 2015; Kaksonen and Roux, 2018), we conclude that α-Syn facilitates endocytosis by enriching the PM with PI4,5P_2_.

In line with previous reports (Nemani et al., 2010; Scott and Roy, 2012; Atias et al., 2019), the results show that WT α-Syn expression inhibits the overall extent of SVs cycling, determined by sypHy signal. This inhibition may result from attenuated mobility of the recycling pool (RP) of SVs in presynaptic boutons and/or reduced pool size at synapses. We interpreted the lower peak level of the sypHy signal, detected in α-Syn expressing neurons, to represent an acceleration in the rate of endocytosis. However, our results also confirm previous observations indicating a reduction in the total secretory capacity of the presynaptic terminals (Nemani et al., 2010; Scott and Roy, 2012; Atias et al., 2019; Sun et al., 2019), which may result from changes in SV pool (Cabin et al., 2002; Chandra et al., 2004), structural and ultrastructural changes in the synapse (Vargas et al., 2017). Overall, the rate of SVs endocytosis in the tested α-Syn forms, represented by the decay in sypHy signal, correlates with their effects on PI4,5P_2_ levels at the synapse. However, the A30P mutation in α-Syn is exceptional in this sense, it increases PIP_2_ levels, yet is ineffective in SV cycling.

There is a general agreement in the field concerning two features of α-Syn protein, its preference for binding acidic phospholipids (Davidson et al., 1998) and its preference for curved membranes, akin the curvature that typifies synaptic vesicles (Antonny, 2011). Thus, the findings herein, indicating that α-Syn interacts with and regulates PI4,5P_2_ levels, fit well with these two features. PI4,5P_2_ is an acidic phospholipid enriched on presynaptic membranes and due to its enrichment with PUFAs, it helps to form membrane curvature (Balla, 2013). PI4,5P_2_ is critical for both mechanisms, endocytosis and exocytosis. Thus, recruiting PIP_2_ to one mechanism will inevitable affect the other. Together with our previous findings, indicating a role for α-Syn to enrich membrane phospholipids with PUFA and increase membrane fluidity (Sharon et al., 2003b), it appears that α-Syn plays major roles in shaping membrane content and in accord, membrane function.

An emerging question in α-Syn’s involvement in mechanisms of membrane trafficking is Why do different studies report a different outcome and how can we get better consistency? To be able to solve this problem we may need to consider the following: 1. The type of endocytic mechanism; 2. Neuronal activity; 3. α-Syn expression; and 4. The lipid content at the plasma membrane. While CME is a key mechanism of SV endocytosis, additional routes of SV endocytosis, including, kiss-and-run, ultrafast and bulk endocytosis take place at the synapse (Gan and Watanabe, 2018; Milosevic, 2018). The degree of involvement and relative importance of each of these mechanisms during physiological neuronal function and in different neuronal types is not fully clear. It is possible that different neurons rely on different mechanisms of endocytosis, depending on their electrophysiological activity and the accompanied need in vesicle recycling (Gan and Watanabe, 2018; Milosevic, 2018). In relevance to neuronal activity, it was suggested that α-Syn role in endocytosis may differ between basal and intense neurotransmission (Lautenschläger et al., 2017). Moreover, neural activity has been shown to control the synaptic accumulation of α-Syn (Fortin, 2005). Thus, the type of α-Syn expression model may affect the outcome, whether α-Syn^−/−^; α, β,γ Syn^−/−^; stable (long term) or transient α-Syn over-expression; exogenously added or endogenously expressed α-Syn. α-Syn is a highly dynamic protein which responds to changes in its environment with structural changes that may affects its activity. Due to its multifaceted nature, it is not possible to consider an opposite outcome when comparing the results obtained in α-Syn silencing vs α-Syn over-expressing models. The results herein indicate that α-Syn actively regulates plasma membrane levels of PI4,5P_2_ to stimulate CME. Considering the above and the additional cellular mechanisms that rely on PI4,5P_2_ levels, it is important to also take PIPs homeostasis into consideration when analyzing α-Syn effects in membrane trafficking.

Abnormal homeostasis of PIPs links defects in membrane trafficking with neurodegeneration (Azarnia Tehran et al., 2018; Raghu et al., 2019). Mutations in Synaptojanin-1 (SynJ1), a PIP-phosphatase enriched in the brain, either in its Sac domain (R258Q and R459P), that dephosphorylates PI4P and PI3P to phosphatidyl inositol (PI) or in the 5-phosphatase domain (Y832C), that dephosphorylates PI4,5P_2_ to PI4P, have been associated with early onset and typical PD (Krebs et al., 2013; Quadri et al., 2013; Kirola et al., 2016; Xie et al., 2019). Mice modeling loss of SynJ1 function, either carrying a PD-causing mutation (R258Q) or by haploinsufficiency (SynJ1 ^+/−^), demonstrate evidence for degeneration of the nigrostriatal dopaminergic system and accumulation of α-Syn pathology (Cao et al., 2017; Pan et al., 2020). The alterations in PIP homeostasis resulting from loss of SynJ1 activity has been associated with impaired autophagy, endocytic disfunction and axonal damage (Cao et al., 2017, 2020; Pan et al., 2020). Of relevance, in a recent report, we linked α-Syn’ physiological activity in PI4,5P_2_ homeostasis with regulation of axonal plasticity and arborization. Importantly, the alterations in PI4,5P_2_ homeostasis were reversed with the expression of SynJ1. We further described evidence for a pathogenic role for α-Syn in dysregulating PI4,5P_2_ in PD (Schechter et al., 2020). Here we extend these findings to show that α-Syn-mediated alterations in PIPs are also involved in its regulation of SV recycling and CME.

## Acknowledgments

The authors thank Prof. Dr. Volker Haucke (Leibniz-FMP Berlin) for critical discussions. This study was funded by Israel Science Foundation grant #182/12 (RS); grant # 1427/12 and #1310/19 (DG).

---

**Supplementary Fig. 1.**
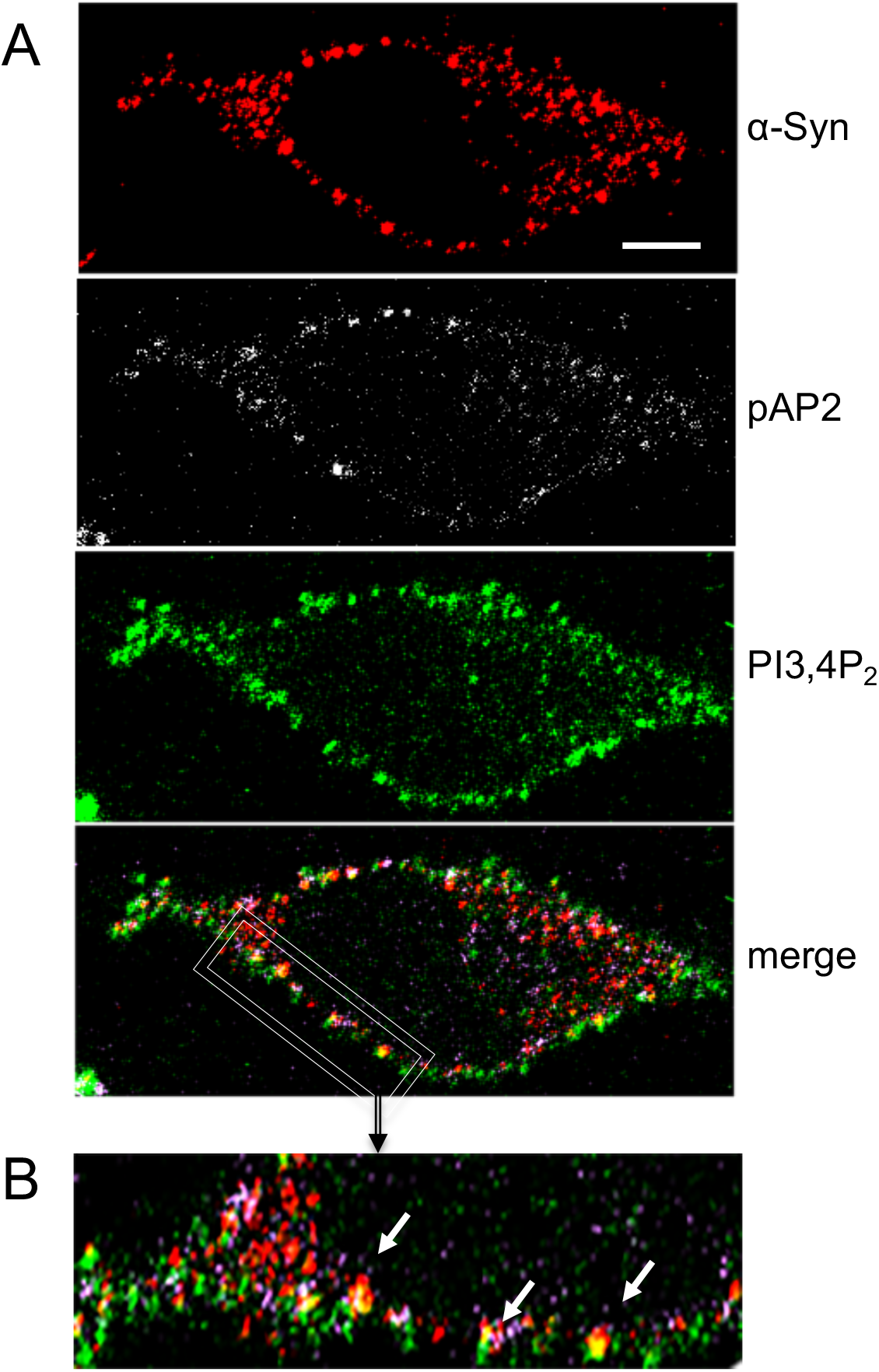
α-Syn colocalizes with phosphorylated AP2 and PIP_2_ on clathrin-coated pits. **A.** SK-Mel2 cells were processed for the detection of the immunoreactive signals of endogenous α-Syn (red), phosphorylated Thr153 μ2 subunit of AP2 (pAP2; Gray) and PI3,4P_2_ (green) by ICC. Bar= 10 μM. **B.** Higher magnification of the image shown in (A), focusing on plasma membrane. Arrows indicate spots of colocalization for α-Syn/pAP2/PI3,4P_2_.

